# DeepKhib: a deep-learning framework for lysine 2-hydroxyisobutyrylation sites prediction

**DOI:** 10.1101/2020.08.14.250712

**Authors:** Luna Zhang, Yang Zou, Ningning He, Yu Chen, Zhen Chen, Lei Li

## Abstract

As a novel type of post-translational modification, lysine 2-Hydroxyisobutyrylation (K_hib_) plays an important role in gene transcription and signal transduction. In order to understand its regulatory mechanism, the essential step is the recognition of K_hib_ sites. Thousands of K_hib_ sites have been experimentally verified across five different species. However, there are only a couple traditional machine-learning algorithms developed to predict K_hi_b sites for limited species, lacking a general prediction algorithm. We constructed a deep-learning algorithm based on convolutional neural network with the one-hot encoding approach, dubbed CNN_OH_. It performs favorably to the traditional machine-learning models and other deep-learning models across different species, in terms of cross-validation and independent test. The area under the ROC curve (AUC) values for CNN_OH_ ranged from 0.82 to 0.87 for different organisms, which is superior to the currently-available K_hib_ predictors. Moreover, we developed the general model based on the integrated data from multiple species and it showed great universality and effectiveness with the AUC values in the range of 0.79 to 0.87. Accordingly, we constructed the on-line prediction tool dubbed DeepKhib for easily identifying K_hib_ sites, which includes both species-specific and general models. DeepKhib is available at http://www.bioinfogo.org/DeepKhib.

## 1 Introduction

Protein post-translational modification (PTM) is a key mechanism to regulate cellular functions through covalent modification and enzyme modification, which dynamically regulates a variety of biological events [1, 2]. Recently, an evolutionarily conserved short-chain lysine acylation modification dubbed lysine 2-hydroxyisobutylation (K_hib_) has been reported, which introduces a steric bulk with a mass shift of +86.03Da (Fig. S1A) and neutralize the positive charge of lysine [3, 4]. It involves various biological functions including biosynthesis of amino acids, starch biosynthesis, carbon metabolism, glycolysis / gluconeogenesis and transcription [3, 5-11]. For instance, the decrease of this modification on K281 of glycolytic enzyme ENO1 reduces its catalytic acitivitie [12]. The three-dimension structure of the peptide containing K281 in the center was shown as Fig. S1B.

Thousands of K_hib_ sites have been identified in different species including humans, plants and prokaryotes through large-scale experimental approaches [3, 5], which is summarized in Table S1. The experimental methods, however, are time-consuming and expensive and thus the development of prediction algorithms in silico is necessary for the high-throughput recognition of K_hib_ sites. Two classifiers (ie. iLys-Khib and Khibpred) have been reported for predicting the K_hib_ sites in a few species [13, 14]. As many different organisms have been investigated and the number of K_hib_ sites has increased, it is indispensable to compare the characteristics of this modification in different species and investigate whether it is suitable to develop a general model with high confidence. Additionally, the reported models were based on traditional machine-learning (ML) algorithms (e.g. Random Forest (RF)). Recently, the deep learning (DL) algorithms, as the modern ML architecture, have demonstrated superior prediction performance in the field of bioinformatics, such as the prediction of modification sites on DNA, RNA and proteins [15-19]. We have developed a few DL approaches for the prediction of PTM sites and they all demonstrate their superiority over conventional ML algorithms [20-22]. Therefore, we attempted to compare the DL models with the traditional ML models for the prediction of K_hib_ sites.

In this study, we constructed a convolutional neural network (CNN)-based architecture with one-hot encoding approach, named as CNN_OH_. This model performed favorably to the traditional ML models and other DL models across different species, in terms of cross-validation and independent test. It is also superior to the documented K_hib_ predictors. Furthermore, we constructed a general model based on the integrated data from multiple species and it demonstrated great generality and effectiveness. Finally, we shared both species-specific models and the general model as the on-line prediction tool DeepKhib for easily identifying K_hib_ sites.

## 2 Materials and Methods

### 2.1 Dataset collection

The experimentally identified K_hib_ sites from five different organisms including *Homo sapiens* (human), *Oryza sativa* (rice), *Physcomitrella patens* (moss) and two one-celled eukaryotes *Toxoplasma gondii* and *Saccharomyces cerevisiae*. The data of the species were pre-processed and the related procedure was exemplified using the human data, as listed below (Fig. S2).

We collected 12,166 K_hib_ sites from 3,055 human proteins [5, 6]. These proteins were classified into 2,466 clusters using CD-HIT with the threshold of 40% according to the previous studies [23, 24]. In each cluster, the protein with the most K_hib_ sites was selected as the representative of the cluster. On the 2,466 representatives, 9,473 K_hib_ sites were considered positives whereas the remaining K sites were taken as negatives. We further estimated the potential redundancy of the positive sites by extracting the peptide segment of seven residues with the K_hib_ site in the center and count the number of unique segments [20, 25]. The number (9,444) of the unique segments is 99.7% of the total segments, suggesting considerable diversity of the positive segments. The number of the negative sites (103,987) is 11 times larger than that of the positive sites. To avoid the potential impact of biased data on model construction, we referred to previous studies and balanced positives and negatives by randomly selecting the same number of negative sites [16, 19]. These positives and negatives composed the whole human dataset.

To determine the optimal sequence window for model construction, we tested different sequence window sizes ranging from 21 to 41, referring to the previous PTM studies where the optimal window sizes are between 31 to 39 [12][17, 20]. The window size of 37 corresponded to the largest area under the ROC curve (AUC) through ten-fold cross-validation (Fig. S3) and was therefore selected in this study. It should be noted that if the central lysine residue is located near the N-terminus or C-terminus of the protein sequence, the symbol “X” is added at the related terminus to ensure the same window size of the sequences.

Fig. 1 showed the flowcharts for all the species. The dataset of each species was randomly separated into five groups of which four were used for ten-fold cross-validation and the rest for independent test. Each group contained the same number of positives and negatives. Specifically, the cross-validation datasets included 15,156/15,464/10,204/12,354 samples for *H. sapiens/T. gondii/O. sativa/P. patens*, respectively. Accordingly, the independent test sets comprised 3,790/3,866/2,552/3,090 samples for these organisms, separately. These datasets are available at http://www.bioinfogo.org/DeepKhib.

**Fig 1.**
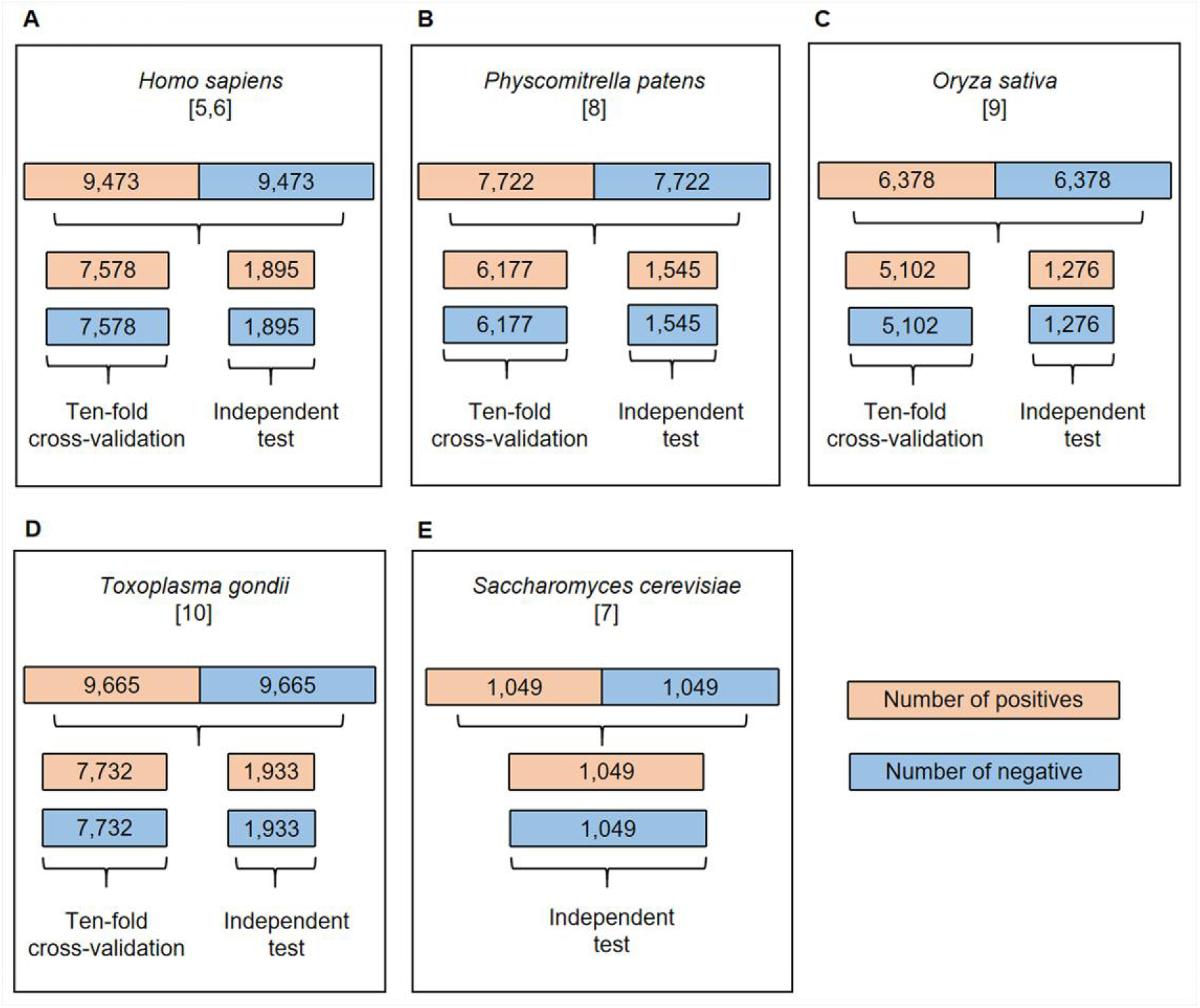
The flowchart of dataset process for *H. sapiens* (A), *P. patens* (B), *O. sativa* (C), *T. gondii* (D) and *S. cerevisiae* (E). All the datasets were separated into cross-validation and independent test datasets except the *S. cerevisiae* dataset.

### 2.2 Feature encodings

#### 2.2.1 The ZSCALE encoding

Each amino acid is characterized by five physiochemical descriptor variables [26, 27].

#### 2.2.2 The encoding of extended amino acid composition (EAAC) encoding

The EAAC encoding is based on the calculation of the amino acid composition (AAC) that indicates the amino acid frequencies for every position in the sequence window. EAAC is calculated by continuously sliding using a fixed-length sequence window (the default is 5) from the N-terminus to the C-terminus of each peptide [28]. The related formula is listed below:

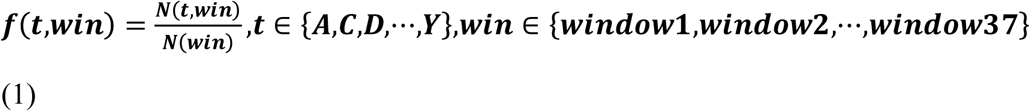

where N (t, win) is the number of amino acid t in the sliding window win, and N(win) is the size of the sliding window win.

#### 2.2.3 The enhanced grouped amino acids content (EGAAC) encoding

The EGAAC feature [22] is developed based on the grouped amino acids content (GAAC) feature [28, 29]. In the GAAC feature, the 20 amino acid types are categorized into five groups (g1: GAVLMI, g2: FYW, g3: KRH, g4: DE and g5: STCPNQ) according to their physicochemical properties and the frequencies of the groups are calculated for every position in the sequence window. For the EGAAC feature, the GAAC values are calculated in the window of fixed length (the default as 5) continuously sliding from the N- to C-terminal of each peptide sequence.

#### 2.2.4 The One-hot encoding

The one-hot encoding is represented by the conversion of the 20 types of amino acids to 20 binary bits. By considering the complemented symbol “X”, a vector of size (20+1) bits is used to represent a single position in the peptide sequence. For example, the amino acid “A” is represented by “100000000000000000000”, “Y” is represented by “000000000000000000010”, and the symbol “X” is represented by “000000000000000000001”.

### 2.3 Architecture of the machine-learning models

#### 2.3.1 The CNN model with one-hot encoding

The CNN algorithm [30] decomposes an overall pattern into many sub-patterns (features) through a neurocognitive machine, and then enters the hierarchically connected feature plane for processing. The architecture of the CNN model with one-hot encoding (called as CNN_OH_) contained four layers as follows (Fig. 2A).

**Fig 2.**
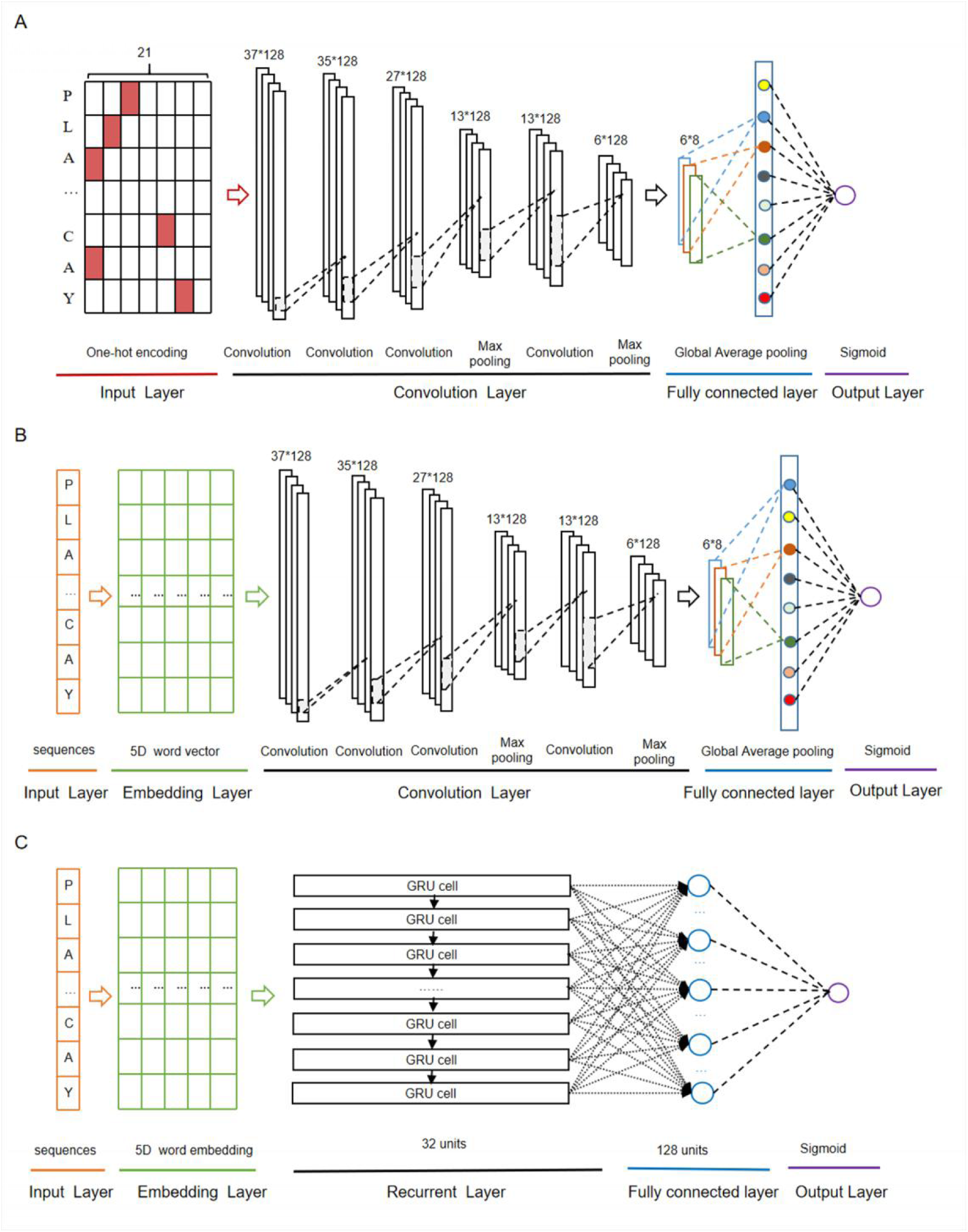
The deep-learning architectures for CNN_OH_ (A), CNN_WE_ (B) and GRU_WE_ (C).

i. The first layer was the input layer where peptide sequences were represented using the one-hot encoding approach.
ii. The second layer was the convolution layer that consisted of four convolution sublayers and two max pooling sublayers. The convolution sublayers, each sublayer uses 128 convolution filters, the length of which are 1, 3, 9 and 10 respectively. The two max pooling sublayers followed the third and fourth convolution sublayers, individually.
iii. The third layer contained the fully connected sublayer, which contained a fully connected sublayer with eight neuron units without flattening, and a global average pooling sublayer, which was adopted to correlate the feature mapping with category output in order to reduce training parameters and avoid over-fitting.
iv. The last layer was the output layer that included a single unit outputting the probability score of the modification, calculated using the “Sigmoid” function. If the probability score is greater than a specified threshold (e.g. 0.5), the peptide is predicted to be positive.

The “ReLU” function [31] was used as the activation function of the convolution sublayers and fully connected sublayers of the above layers to avoid gradient dispersion in the training process. The Adam optimizer [32] was used to optimize the hyper-parameters of this model, which include batch size, maximum epoch, learning rate and dropout rate. The maximum training period was set as 1000 epochs to ensure the convergence of the loss function values. In each epoch, the training data set was separated and iterated in a batch size of 1024. To avoid over-fitting, the dropout of neurons units in each convolution sublayer of the second layer was set 70% and that in the full connection sublayer of the third layer was set 30% [33], the early stop strategy was adopted and the best model was saved.

#### 2.3.2 The CNN algorithm with word embedding

The CNN algorithm with word embedding (CNN_WE_) contained five layers (Fig. 2B). The input layer receives the sequence of window size 37 and each residue is transformed into a five-dimensional word vector in the embedding layer. The rest layers are the same as the corresponding layers in CNN_OH_.

#### 2.3.3 The GRU algorithm with word embedding

The GRU algorithm [34] includes an update gate and a reset gate. The former is used to control the extent to which the state information at the previous moment is brought into the current state, whereas the latter is used to control the extent to which the state information at the previous moment is ignored. The GRU algorithm with word embedding (GRU_WE_) contained five layers (Fig. 2C). The first, the second and the last layers are the same as the corresponding layers in CNN_WE_. The third layer is the recurrent layer where each word vector from the previous layer was sequentially inputted into the related GRU unit that contains 32 hidden neuron units. The fourth layer was the fully connected layer that contains 128 neuron units with “ReLU” as the activation function.

#### 2.3.4 The RF algorithms with different features

The Random Forest algorithm [35] contains multiple decision trees, which remain unchanged under the scaling of feature values and various other transformations, and the output category is determined by the mode of the category output by the individual tree. The RF algorithm integrates multiple decision trees and chooses the classification with the most votes from the trees. Each tree depends on the values of a random vector sampled independently with the same distribution for all trees in the forest. The number of decision trees was set 140. This classifier was developed based on the Python module “sklearn”.

### 2.4 Cross-validation and Performance evaluation

To evaluate the performance of K_hib_ sites prediction, we adopted four statistical measurement methods. They included sensitivity (Sn), specificity (Sp), accuracy (ACC), and Matthew’s correlation coefficient (MCC), listed as follows:

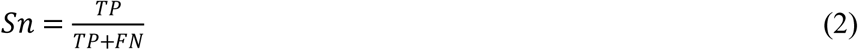

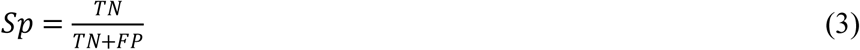

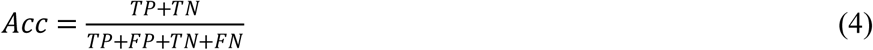

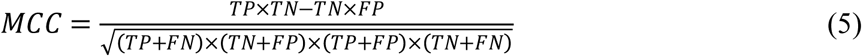

In the above equations, TP is true positives, FP is false positives, TN is true negatives, FN is false negatives. In addition, the area under the receiver operating characteristic (ROC) curve (AUC) values was calculated to evaluate the performance of the prediction model.

### 2.5 Statistical methods

The paired student’s t-test was used to test the significant difference between the mean values of the two paired populations. As for multiple comparisons, the adjusted P value with the Benjamini-Hochberg (BH) method was adopted.

## 3 Results and discussion

A couple of computational approaches has been developed for the prediction of K_hib_ sites [13, 14]. Recently, this modification has been investigated across five different species, ranging from single-celled organisms to multiple-celled organisms and from plants to mammals. Additionally, the number of reported sites has been significantly increased. These raised our interest to develop novel prediction algorithms and explore the characteristics of this modification. We pre-processed the data from different species and separated them into the cross-validation dataset and the independent test set (see Methods for detail; Fig. 1). We first took the human data as the representative to compare different models and then applied the model with the best performance to other species. The human cross-validation dataset contained 15,156 samples and the independent test set covered 3,790 samples, in each of which half were positives and half were negatives.

### 3.1 CNN_OH_ showed superior performance

We constructed nine models, divided into two categories: six traditional ML models and three DL models (See Methods for details). The traditional ML models were based on the RF algorithm combined with different encoding schemes. The DL models included a Gated Recurrent Unit (GRU) model with the word-embedding encoding approach dubbed GRU_WE_ and two CNN models with the one-hot and word-embedding encoding approaches named CNN_OH_ and CNN_WE_, respectively. Both encoding methods are common in the DL algorithms [20, 25].

The RF-based models were developed with different common encoding schemes, including EAAC, EGAAC and ZSCALE. Among these encoding schemes, EGAAC had the best performance followed by EAAC whereas ZSCALE was the worst in terms of AUC and ACC for both ten-fold cross-validation and the independent test (Table 1, Fig. 3). For instance, EGAAC corresponded to the average AUC value as 0.775, EAAC had the value as 0.763 and ZSCALE had the value as 0.740 for cross validation. Because different encodings represent distinct characteristics of K_hib_-containing peptides, we evaluated the combinations of the encoding schemes. The combinations showed better performances than individual scheme and the combination of all the three was the best for both cross-validation and the independent test, in terms of AUC, MCC and ACC (Table 1, Fig. 3). Therefore, the K_hib_ prediction accuracy could be improved by the integration of different encoding schemes.

**Table 1.** Performances comparison of the different classifiers for human K_hib_ prediction.

|  | Classifier | Sn | Sp | Acc | MCC | AUC |
| --- | --- | --- | --- | --- | --- | --- |
| Ten-fold<br>cross-validation | RF<sub>EGAAC</sub> | 0.727±0.015 | 0.682±0.017 | 0.704±0.011 | 0.409±0.022 | 0.775±0.011 |
|  | RF<sub>EAAC</sub> | 0.744±0.025 | 0.645±0.023 | 0.695±0.010 | 0.391±0.020 | 0.763±0.008 |
|  | RF<sub>ZSCALE</sub> | 0.681±0.016 | 0.662±0.018 | 0.672±0.011 | 0.344±0.023 | 0.740±0.014 |
|  | RF<sub>EGAAC+EAAC</sub> | 0.748±0.019 | 0.691±0.023 | 0.719±0.012 | 0.439±0.025 | 0.789±0.011 |
|  | RF<sub>EGAAC+ZSCALE</sub> | 0.726±0.019 | 0.707±0.015 | 0.716±0.012 | 0.433±0.025 | 0.794±0.010 |
|  | RF<sub>EGAAC+EAAC+ZSCALE</sub> | 0.751±0.016 | 0.702±0.022 | 0.727±0.013 | 0.454±0.026 | 0.802±0.010 |
|  | GRU<sub>WE</sub> | 0.821±0.024 | 0.683±0.033 | 0.752±0.009 | 0.509±0.018 | 0.830±0.007 |
|  | CNN<sub>WE</sub> | 0.849±0.035 | 0.722±0.042 | 0.786±0.007 | 0.578±0.012 | 0.867±0.005 |
|  | CNN<sub>OH</sub> | 0.876±0.025 | 0.700±0.026 | 0.788±0.007 | 0.586±0.014 | 0.868±0.004 |
| Independent test | RF<sub>EGAAC</sub> | 0.719±0.006 | 0.676±0.007 | 0.698±0.002 | 0.395±0.004 | 0.767±0.002 |
|  | RF<sub>EAAC</sub> | 0.755±0.003 | 0.638±0.007 | 0.697±0.003 | 0.396±0.006 | 0.764±0.003 |
|  | RF<sub>ZSCALE</sub> | 0.680±0.008 | 0.658±0.009 | 0.669±0.005 | 0.337±0.011 | 0.736±0.003 |
|  | RF<sub>EGAAC+EAAC</sub> | 0.740±0.006 | 0.678±0.005 | 0.709±0.002 | 0.419±0.005 | 0.781±0.002 |
|  | RF<sub>EGAAC+ZSCALE</sub> | 0.728±0.006 | 0.692±0.006 | 0.710±0.002 | 0.420±0.005 | 0.787±0.002 |
|  | RF<sub>EGAAC+EAAC+ZSCALE</sub> | 0.752±0.005 | 0.693±0.004 | 0.723±0.002 | 0.446±0.005 | 0.796±0.002 |
|  | GRU<sub>WE</sub> | 0.806±0.015 | 0.692±0.029 | 0.749±0.004 | 0.501±0.007 | 0.824±0.005 |
|  | CNN<sub>WE</sub> | 0.846±0.035 | 0.719±0.042 | 0.783±0.006 | 0.572±0.009 | 0.865±0.004 |
|  | CNN<sub>OH</sub> | 0.874±0.026 | 0.690±0.035 | 0.782±0.005 | 0.575±0.005 | 0.871±0.001 |

Note: The data sets for ten-fold cross-validation and an independent test were described in the Methods. The RF classifier with the different encoding approach was named as RF<sub>EGAAC</sub>, RF<sub>EAAC</sub>, RF<sub>ZSCALE</sub>, RF<sub>EGAAC+EAAC</sub>, RF<sub>EGAAC+ZSCALE</sub> and RF<sub>EGAAC+EAAC+ZSCALE</sub>. The RNN/CNN classifier with the word embedding encoding approach was named as GRU<sub>WE</sub>/CNN<sub>WE</sub>, respectively. The CNN classifier with one-hot encoding was named as CNN<sub>OH</sub>. Ten models were constructed in the ten-fold cross validation and evaluated using the ten different validation datasets and the same independent dataset. Accordingly, the value Sn, Sp, Acc, MCC and AUC were represented by average ±standard deviation.

**Fig 3.**
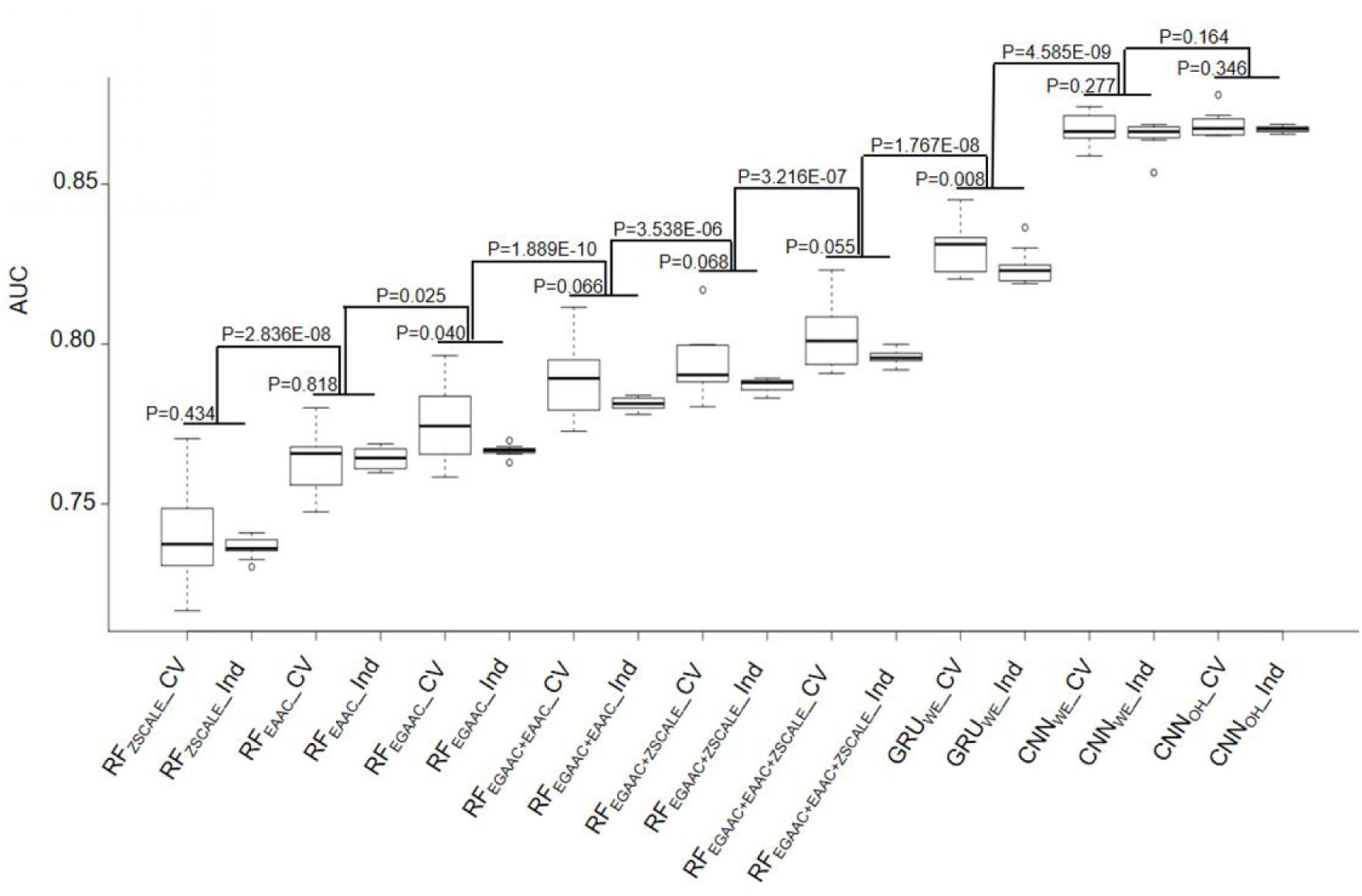
Performance comparison of ten-fold cross-validation and independent test datasets of nine different models.

As the DL algorithms showed superior to the traditional ML algorithms for a few PTM predictions in our previous studies [21, 22], we examined the DL algorithms for the K_hib_ prediction. Traditionally, CNN is popular for image prediction with spatial invariant features while RNN is ideal for text prediction with sequence features. However, many cases demonstrate that CNN also has good performance when applied to sequence data [16, 36]. Accordingly, we developed both RNN and CNN models for the K_hib_ prediction with two common encoding approaches: one-hot and word-embedding. Expectedly, all three DL models were significantly better than the traditional ML models constructed above in the cross-validation and independent test (Table 1, Fig. 3). For instance, the average AUC values of the DL models were above 0.824 whereas those of the ML models were less than 0.802.

In these DL models, two CNN models CNN_OH_ and CNN_WE_ had similar performances and both compared favorably to GRU_WE_ (Table 1, Fig. 3). CNN_OH_ had the AUC value as 0.868 for the cross-validation and its values of SN, SP, ACC and MCC were 0.876, 0.700, 0.788 and 0.586, respectively. Here, we chose CNN_OH_ as the 2-Hydroxyisobutyrylation predictor. We evaluated the robustness of our models by comparing their performances between the cross-validation and independent tests. As their performances between these two tests had no statistically different (P>0.01), we concluded that our constructed models were robust and neither over-fitting nor under-fitting.

### 3.2 Construction and comparison of predictors for other species

We constructed nine models for the human organism and chose CNN_OH_ as the final prediction model. We applied the CNN_OH_ architecture to the other three organisms (i.e. *T. gondii, O. sativa and P. patens*). For each organism, we separated the dataset as the cross-validation set and the independent set. Similar to the human species, the CNN_OH_ models for these species had similar performances between cross-validation and independent test and their AUC values were larger than 0.818 (Table 2). It indicates that these constructed models are effective and robust.

**Table 2.** The AUC values of the CNN_OH_ model constructed for *O. sativa, P. patens, T. gondii*, and *H. sapiens*, respectively.

| Species | Ten-fold cross-validation | Independent test |
| --- | --- | --- |
| *O. sativa* | 0.823 | 0.818 |
| *P. patens* | 0.830 | 0.831 |
| *T. gondii* | 0.862 | 0.865 |
| *H. sapiens* | 0.868 | 0.871 |

As lysine 2-Hydroxyisobutyrylation is conserved across different types of species, we hypothesized that the model built for one species may be used to predict K_hib_ sites for other species. To test this hypothesis, we compared the performances of the CNN_OH_ models in terms of the independent data sets of individual species. Additionally, we built a general CNN_OH_ model based on the training datasets integrated from all the four species. Table 3 shows that the AUC values of these predictions were larger than 0.761, suggesting that the cross-species prediction had reliable performances. Specifically, given a species, the best prediction performances were derived from the general model and the model developed specifically for this species. For instance, the human CNN_OH_ model had the best performance followed by the general model in terms of the human independent test whereas the general model had the best accuracy followed by the moss-specific model for the moss independent test. These suggest that on one hand, lysine 2-Hydroxyisobutyrylation of each species has its own characteristics; one the other hand, this modifications across different species share strong commonalities. Therefore, the general model may be effectually applied to any species. Furthermore, we evaluated the generality of the general CNN_OH_ model using the dataset of *S. cerevisiae* that contained 1,049 positive and 1,049 negative samples, which may not be enough for build an effective DL predictor [20]. The general model got the AUC value as 0.789, indicating the generality of this model. In other words, the general model is effective to predict K_hib_ sites for any organism.

**Table 3.** The AUC values of different CNN_OH_ models in terms of independent test for five distinct organisms.

| Prediction models | Independent data sets |  |  |  |  |
| --- | --- | --- | --- | --- | --- |
|  | *O. sativa* | *P. patens* | *T. gondii* | *H. sapiens* | *S. cerevisiae* |
| *O. sativa* | **0.818** | 0.788 | 0.782 | 0.803 | 0.721 |
| *P. patens* | 0.761 | **0.831** | 0.812 | 0.837 | **0.806** |
| *T. gondii* | 0.781 | 0.813 | **0.865** | 0.827 | 0.776 |
| *H. sapiens* | 0.778 | 0.818 | 0.832 | **0.871** | 0.785 |
| General | **0.802** | **0.840** | **0.860** | **0.868** | **0.789** |

Note: The top two models with best performance are bold.

We identified and compared the significant patterns and conserved motifs between K_hib_ and non-K_hib_ sequences across the different organisms using the two-sample-logo program with t-test (P<0.05) with Bonferroni correction[37]. Fig. 4 shows the similarities and differences between the species. For instance, the residues R and K at the −1 position (i.e. R&K@P-1) and P at +1 position (i.e. P@P+1) are significantly depleted across the species. On the contrary, K&R@P+1 tend to be enriched for *H. sapiens* but depleted for *T. gondii* whereas both species have the depleted residue Serine across the positions ranging from P-18 to P+18. These similarities between the organisms may result in the generality and effectiveness of the general CNN_OH_ model.

**Fig 4.**
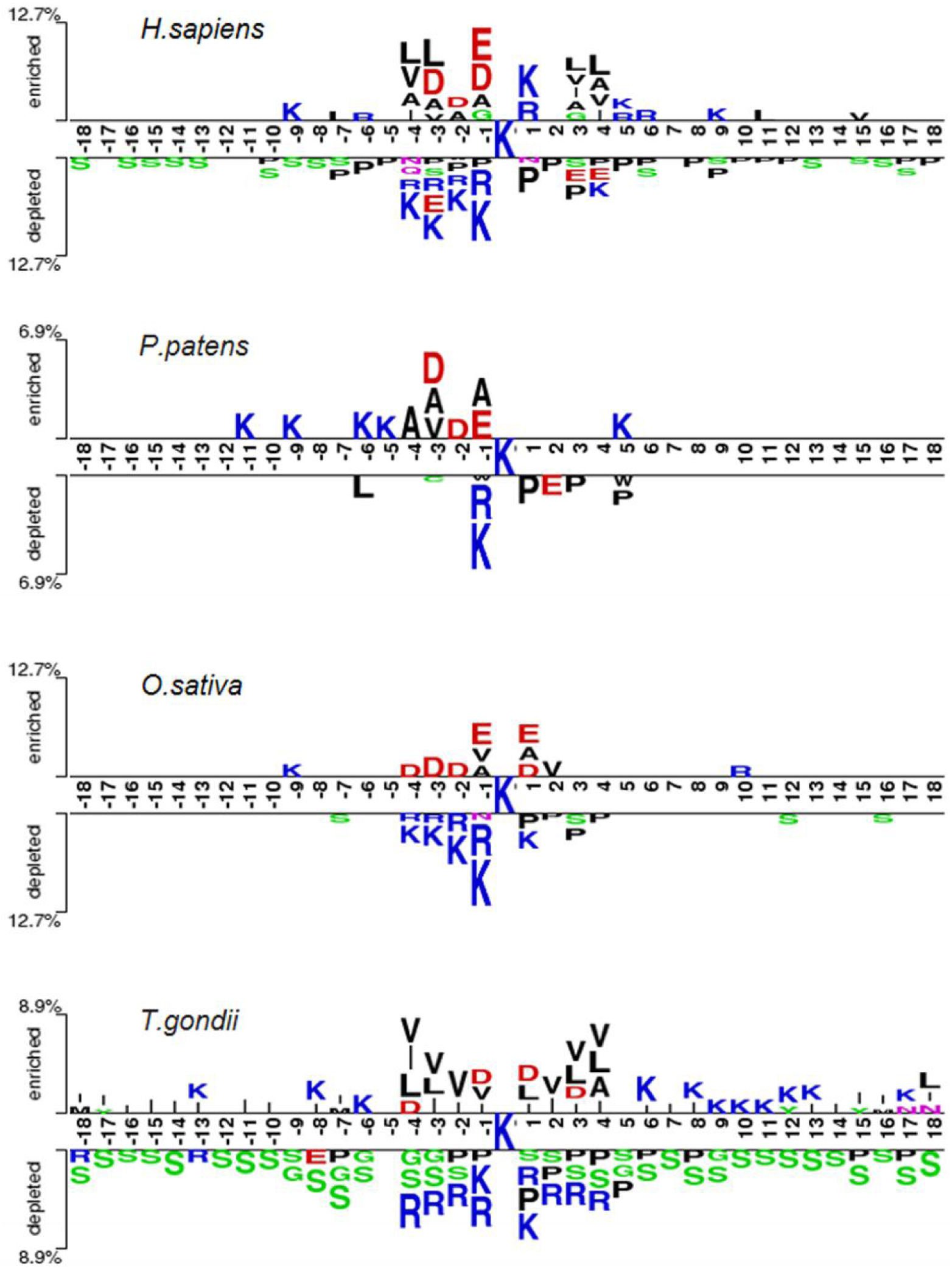
Sequence pattern surrounding the K_hib_ sites, including the significantly enriched and depleted residues based on K_hib_ peptides and non-modification peptides from different species (P<0.05, student’s T-test with Bonferroni correction). The pattern was generated using the two-sample-logo method [37].

### 3.3 Comparison of CNNOH with the reported predictors

We assessed the performance of CNN_OH_ by comparing it with the existing K_hib_ predictors KhibPred[14] and iLys-Khib[13]. First, we compared CNN_OH_ with KhibPred for individual species in terms of ten-fold cross-validation[14]. The average AUC values of CNN_OH_ were 0.868/0.830/0.823 for *H. sapiens/P. patens/O. sativa*, respectively (Table 2). On the contrary, the corresponding values of KhibPred were 0.831/0.781/0.825[14]. Thus, CNN_OH_ compares favorably to KhibPred. Second, the model iLys-Khib was constructed and tested using 9,318 human samples as the ten-fold cross-validation data set and 4,219 human samples as the independent test set. We used the same datasets to construct CNN_OH_ and compared it with iLys-Khib. CNN_OH_ outperformed iLys-Khib in terms of all the measurements of performance (e.g. Sn, Sp, Acc, MCC and AUC) for both ten-fold cross-validation and independent test (Table 4). For instance, the AUC value of CNN_OH_ was 0.860 for the independent test whereas that of iLys-Khib was 0.756. In summary, CNN_OH_ is a competitive predictor.

**Table 4.** The prediction performance of CNN_OH_ compared to iLys-Khib in terms of the same cross-validation and independent test datasets.

| Dataset | Model | Sn | Sp | Acc | MCC | AUC |
| --- | --- | --- | --- | --- | --- | --- |
| Ten-fold cross-validation | iLys-Khib | 0.745 | 0.658 | 0.701 | 0.404 | 0.770 |
|  | CNN<sub>OH</sub> | 0.830 | 0.713 | 0.772 | 0.547 | 0.847 |
| Independent test | iLys-Khib | 0.725 | 0.643 | 0.648 | 0.186 | 0.756 |
|  | CNN<sub>OH</sub> | 0.861 | 0.685 | 0.696 | 0.281 | 0.860 |

### 3.4 Construction of the on-line K_hib_ predictor

We developed an easy-to-use Web tool for the prediction of K_hib_ sites, dubbed as DeepKhib. It contains five CNN_OH_ models, including one general model and four models specific to the species (i.e. *H. sapiens, O. sativa, P. patens and T. gondii*). Given a species of interest, users could select the suitable model (e.g. the general model or the model specific to an organism) for prediction (Fig. 5A). After the protein sequences as the fasta file format are uploaded, the prediction results will be shown with five columns: Protein, Position, Sequence, Prediction score and Prediction category (Fig. 5B). The prediction category covered four types according to the prediction scores: no (0-0.320), medium confidence (0.320-0.441), high confidence (0.441-0.643) and very high confidence (0.643-1).

**Fig 5.**
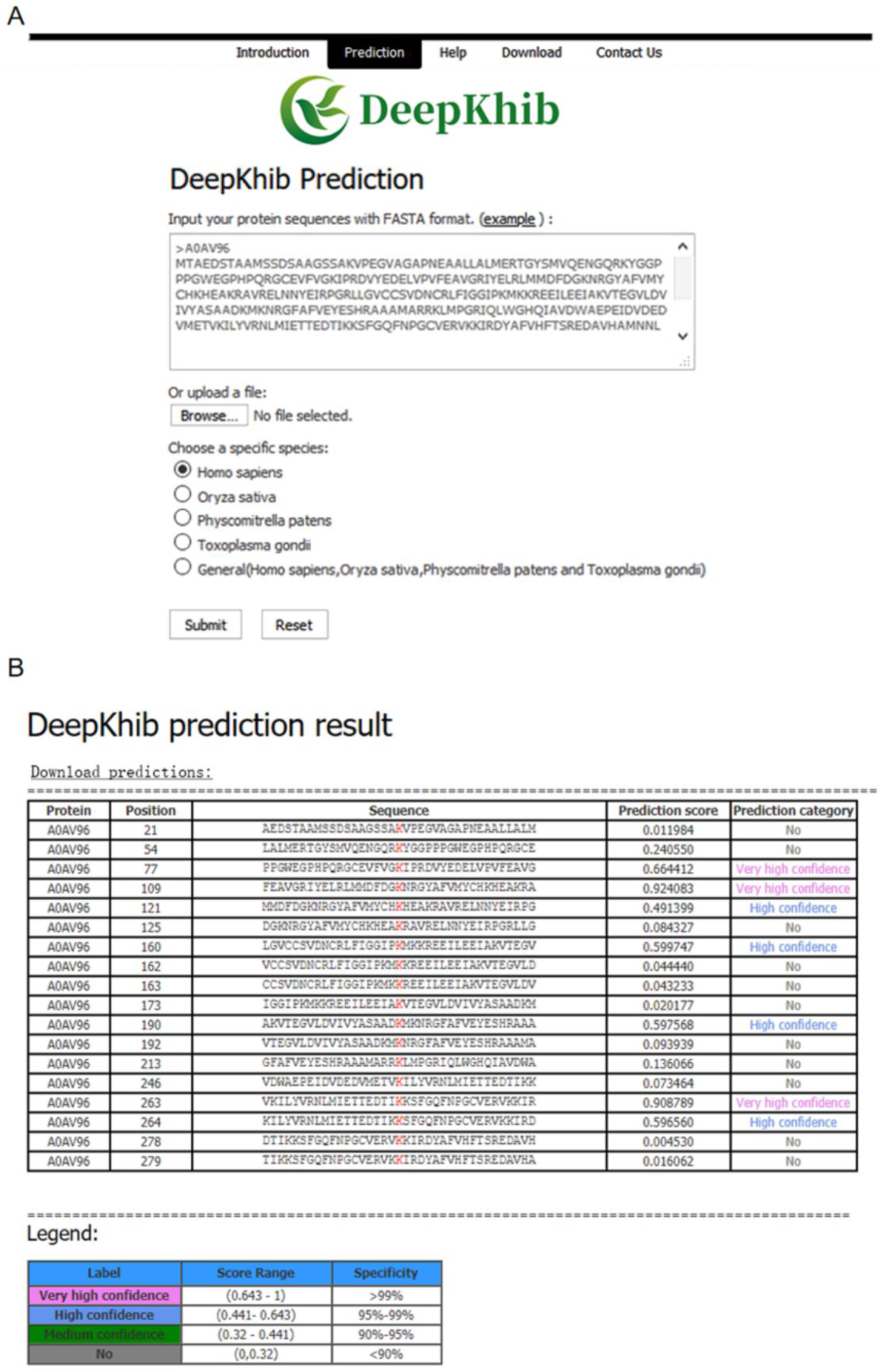
DeepKhib interface for the prediction of K_hib_ sites with the option of organism-specific or general classifiers (A) and its application to the prediction (B).

## 4 Conclusions

The common PTM classifiers are mainly based on the traditional ML algorithms that require the pre-defined informative features. Here, we applied the advanced DL algorithm CNN_OH_ for predicting K_hib_ sites. CNN_OH_ shows its superior performance, because of the capability of the multi-layer CNN algorithm to extract complex features and learn sparse representation in a self-taught manner. Moreover, the general CNN_OH_ model demonstrates great generality and effectiveness, due to the conservation of K_hib_ modification from single-cell to multiple-cell organisms. The outstanding performance of DL in the prediction of K_hib_ sites suggests that DL may be applied broadly to predicting other types of modification sites.

## Authors’ contributions

LL conceived this project. LZ and YZ constructed the algorithms under the supervision of LL and ZC; LZ and NH analyzed the data. LL, YZ, YC and LZ wrote the manuscript. All authors read and approved the final manuscript.

## Acknowledgments

This work was supported in part by funds from the Young Scientists Fund of the National Natural Science Foundation of China (Grant No. 31701142 to ZC), and the National Natural Science Foundation of China (Grant No. 31770821 to LL); LL is supported by the “Distinguished Expert of Overseas Tai Shan Scholar” program. YZ is supported by the Qingdao Applied Research Project.

## References

1. Beltrao, P., et al., Evolution and functional cross-talk of protein post-translational modifications. Mol Syst Biol, 2013. 9: p. 714.

2. Skelly, M.J., L. Frungillo, and S.H. Spoel, Transcriptional regulation by complex interplay between post-translational modifications. Current Opinion in Plant Biology, 2016. 33: p. 126–132.

3. Dai, L., et al., Lysine 2-hydroxyisobutyrylation is a widely distributed active histone mark. Nature Chemical Biology, 2014. 10(5): p. 365–70.

4. Xiao, H., et al., Genetic Incorporation of epsilon-N-2-Hydroxyisobutyryl-lysine into Recombinant Histones. ACS Chem Biol, 2015. 10(7): p. 1599–603.

5. Huang, H., et al., Landscape of the regulatory elements for lysine 2-hydroxyisobutyrylation pathway. Cell Res, 2018. 28(1): p. 111–125.

6. Wu, Q., et al., Global Analysis of Lysine 2-Hydroxyisobutyrylome upon SAHA Treatment and Its Relationship with Acetylation and Crotonylation. J Proteome Res, 2018. 17(9): p. 3176–3183.

7. Huang, J., et al., 2-hydroxyisobutyrylation on histone h4k8 is regulated by glucose homeostasis in saccharomyces cerevisiae. Proceedings of the National Academy of Sciences, 2017. 114(33).

8. Yu, Z., et al., Proteome-wide identification of lysine 2-hydroxyisobutyrylation reveals conserved and novel histone modifications in Physcomitrella patens. Sci Rep, 2017. 7(1): p. 15553.

9. Meng, X., et al., Proteome-wide Analysis of Lysine 2-hydroxyisobutyrylation in Developing Rice (Oryza sativa) Seeds. Sci Rep, 2017. 7(1): p. 17486.

10. Yin, D., et al., Global Lysine Crotonylation and 2-Hydroxyisobutyrylation in Phenotypically Different Toxoplasma gondii Parasites. Molecular & Cellular Proteomics, 2019.

11. Li, Q.Q., et al., Proteomic analysis of proteome and histone post-translational modifications in heat shock protein 90 inhibition-mediated bladder cancer therapeutics. Sci Rep, 2017. 7(1): p. 201.

12. Huang, H., et al., p300-Mediated Lysine 2-Hydroxyisobutyrylation Regulates Glycolysis. Mol Cell, 2018. 70(4): p. 663–678 e6.

13. Ju, Z. and S.-Y. Wang, iLys-Khib: Identify lysine 2-Hydroxyisobutyrylation sites using mRMR feature selection and fuzzy SVM algorithm. Chemometrics and Intelligent Laboratory Systems, 2019. 191: p. 96–102.

14. Wang. YG, et al., Accurate prediction of species-specific 2-hydroxyisobutyrylation sites based on machine learning frameworks. Analytical biochemistry, 2020. 602: p. 113793.

15. Tian, Q., et al., MRCNN: a deep learning model for regression of genome-wide DNA methylation. BMC Genomics, 2019. 20(Suppl 2): p. 192.

16. Tahir, M., H. Tayara, and K.T. Chong, iPseU-CNN: Identifying RNA Pseudouridine Sites Using Convolutional Neural Networks. Mol Ther Nucleic Acids, 2019. 16: p. 463–470.

17. Wang, D., et al., Musitedeep: a deep-learning framework for general and kinase-specific phosphorylation site prediction. Bioinformatics, 2017. 10.

18. Long, H., et al., A Hybrid Deep Learning Model for Predicting Protein Hydroxylation Sites. Int J Mol Sci, 2018. 19(9).

19. Huang, Y., et al., BERMP: a cross-species classifier for predicting mA sites by integrating a deep learning algorithm and a random forest approach. International journal of biological sciences, 2018. 14(12): p. 1669–1677.

20. Chen, Z., et al., Integration of A Deep Learning Classifier with A Random Forest Approach for Predicting Malonylation Sites. Genomics Proteomics Bioinformatics, 2018. 16(6): p. 451–459.

21. Chen. Z, et al., Large-scale comparative assessment of computational predictors for lysine post-translational modification sites. Briefings in bioinformatics, 2019. 20(6): p. 2267–2290.

22. Zhao, Y., et al., Identification of Protein Lysine Crotonylation Sites by a Deep Learning Framework With Convolutional Neural Networks. IEEE Access, 2020. 8: p. 14244–14252.

23. Huang, Y., et al., CD-HIT Suite: a web server for clustering and comparing biological sequences. Bioinformatics, 2010. 26(5): p. 680–2.

24. Li, W. and A. Godzik, Cd-hit: a fast program for clustering and comparing large sets of protein or nucleotide sequences. Bioinformatics, 2006. 22(13): p. 1658–9.

25. Xie, Y., et al., DeepNitro: Prediction of Protein Nitration and Nitrosylation Sites by Deep Learning. Genomics Proteomics Bioinformatics, 2018. 16(4): p. 294–306.

26. Sandberg, M., L. Eriksson, and J. Jonsson, New chemical descriptors relevant for the design of biologically active peptides. a multivariate characterization of 87 amino acids. Journal of Medicinal Chemistry, 1998. 41(14): p. 2481–2491.

27. Chen, Y.Z., et al., Sumohydro: a novel method for the prediction of sumoylation sites based on hydrophobic properties. PLoS ONE, 2012. 7(6).

28. Chen, Z., et al., iFeature: a python package and web server for features extraction and selection from protein and peptide sequences. Bioinformatics, 2018.

29. Chen. Z, et al., iLearn: an integrated platform and meta-learner for feature engineering, machine-learning analysis and modeling of DNA, RNA and protein sequence data. Briefings in bioinformatics, 2020. 21(3): p. 1047–1057.

30. Fukushima, K., Neocognitron: a self organizing neural network model for a mechanism of pattern recognition unaffected by shift in position. Biol Cybern, 1980. 36(4): p. 193–202.

31. Hahnloser, R.H., et al., Digital selection and analogue amplification coexist in a cortex-inspired silicon circuit. Nature, 2000. 405(6789): p. 947–51.

32. Kingma, D.P. and B. J, Adam: A Method for Stochastic Optimization. Computer Science, 2014.

33. Nitish, S., et al., Dropout: a simple way to prevent neural networks from overfitting. 2014. 15: p. 1929–1958.

34. Cho, K., et al., Learning Phrase Representations using RNN Encoder-Decoder for Statistical Machine Translation. Computer Ence, 2014.

35. Breiman, L., Random Forests. Machine Learning, 2001. 45(1): p. 5–32.

36. Sainath, T.N., et al., Deep convolutional neural networks for LVCSR. IEEE International Conference on Acoustic, 2013.

37. Vacic. V, Iakoucheva. LM, and Radivojac. P, Two Sample Logo: a graphical representation of the differences between two sets of sequence alignments. Bioinformatics (Oxford, England), 2006. 22(12): p. 1536–7.

