## Supplementary_Material for "DeepKhib: a deep-learning framework for lysine 2-hydroxyisobutyrylation sites prediction"

Table S1. Lysine 2-hydroxyisobutyrylation in different species.

| **Years** | **Species** | **Proteins** | **Sites** | **Reference** |
| --- | --- | --- | --- | --- |
| 2017 | *Homo sapiens (HeLa cells)* | 1, 725 | 6,548 | [1] |
| 2018 | *Homo sapiens (A549 cells)* | 2,484 | 8,765 | [2] |
| 2017 | *Saccharomyces cerevisiae* | 369 | 1,458 | [3] |
| 2017 | *Physcomitrella patens* | 3,001 | 11,976 | [4] |
| 2017 | *Oryza sativa* | 2,512 | 9,916 | [5] |
| 2019 | *Toxoplasma gondii (RH)；*  *Toxoplasma gondii (ME49)* | 1,950;  1,720 | 9,502;  8,092 | [6] |

A

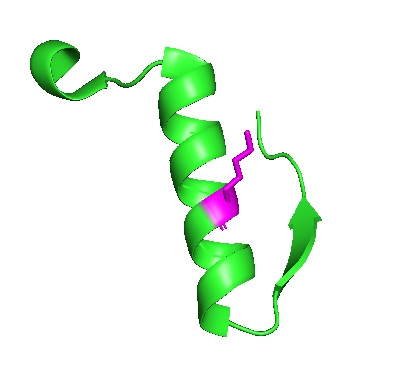

B

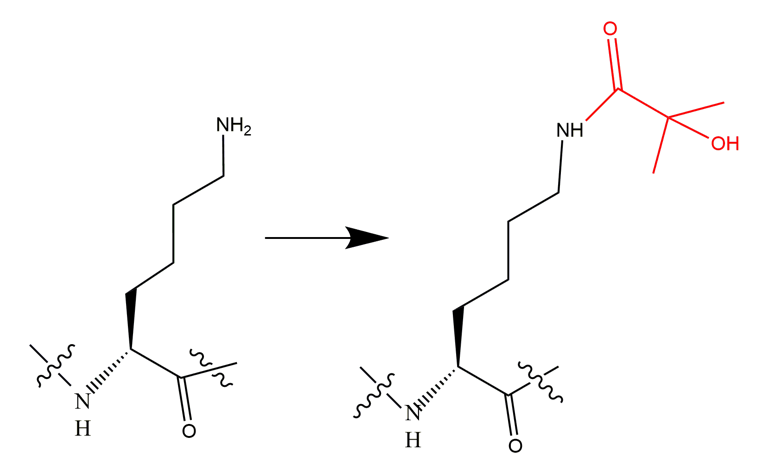

Fig S1. A. Three-dimensional structure of the peptide (15 amino acid long) with K281 in the center that can be hydroxyisobutyrylated from the protein Enolase 1 (from PDB ID: 2PSN)[7]. B. The chemical equation from Lysine to hydroxysobutyrylated lysine.

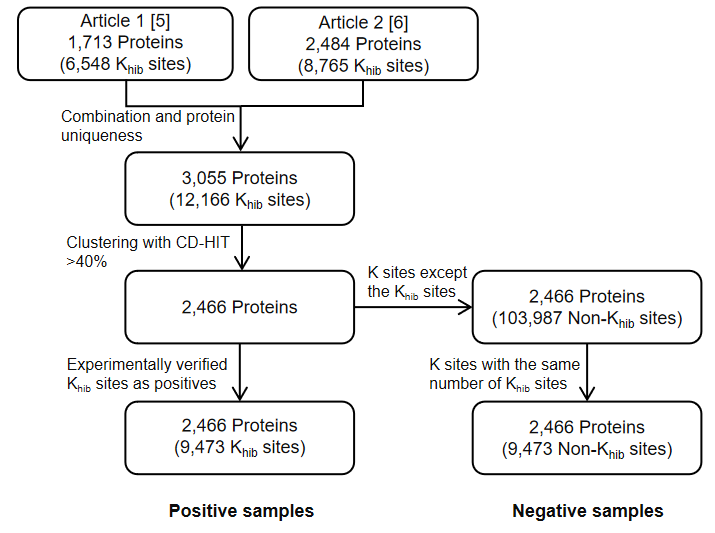

Fig S2. The workflow of data collection and pre-processing for the human dataset.

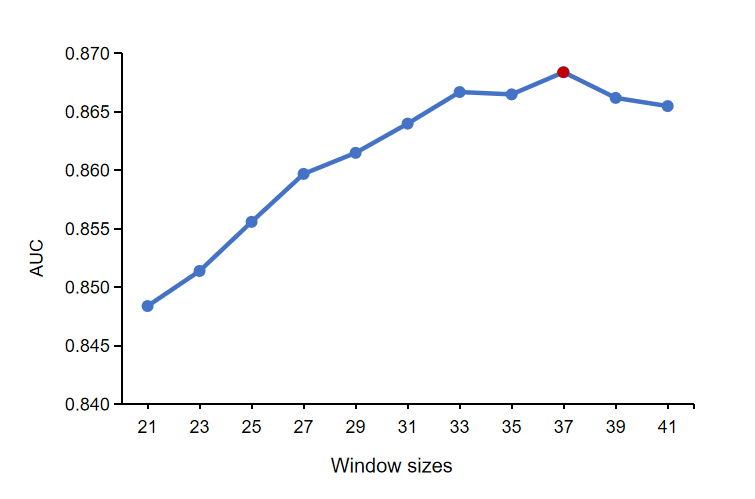

Fig S3. The performance of the CNN_OH_ classifier constructed using different window sizes through the ten-fold cross-validation. Window size of 37 highlighted by red spot was selected as the peptide length for the classifier construction in this study.

**References**

1.	Huang, H., et al., *Landscape of the regulatory elements for lysine 2-hydroxyisobutyrylation pathway.* Cell Res, 2018. **28**(1): p. 111-125.

2.	Wu, Q., et al., *Global Analysis of Lysine 2-Hydroxyisobutyrylome upon SAHA Treatment and Its Relationship with Acetylation and Crotonylation.* J Proteome Res, 2018. **17**(9): p. 3176-3183.

3.	Huang, J., et al., *2-hydroxyisobutyrylation on histone h4k8 is regulated by glucose homeostasis in saccharomyces cerevisiae.* Proceedings of the National Academy of Sciences, 2017. **114**(33).

4.	Yu, Z., et al., *Proteome-wide identification of lysine 2-hydroxyisobutyrylation reveals conserved and novel histone modifications in Physcomitrella patens.* Sci Rep, 2017. **7**(1): p. 15553.

5.	Meng, X., et al., *Proteome-wide Analysis of Lysine 2-hydroxyisobutyrylation in Developing Rice (Oryza sativa) Seeds.* Sci Rep, 2017. **7**(1): p. 17486.

6.	Yin, D., et al., *Global Lysine Crotonylation and 2- Hydroxyisobutyrylation in Phenotypically Different Toxoplasma gondii Parasites.* Molecular & Cellular Proteomics, 2019.

7.	Huang, H., et al., *p300-Mediated Lysine 2-Hydroxyisobutyrylation Regulates Glycolysis.* Mol Cell, 2018. **70**(4): p. 663-678 e6.
